# Overturning the paradigm that IL6 signaling drives liver regrowth while shining light on a new therapeutic target for regenerative medicine

**DOI:** 10.1101/2021.04.05.438446

**Authors:** Jinrui Dong, Sivakumar Viswanathan, Eleonora Adami, Sebastian Schafer, Fathima F. Kuthubudeen, Anissa A. Widjaja, Stuart A. Cook

## Abstract

It is accepted that IL6 signaling in hepatocytes, mediated via glycoprotein 130 (gp130), is beneficial and that HyperIL6 promotes liver regeneration by activating STAT3. Recently, autocrine IL11 activity, which also signals via gp130 and ERK, was found to be hepatotoxic. Here we examined whether the beneficial effects of HyperIL6 could reflect unappreciated competitive inhibition of IL11 signaling. In hepatocytes, HyperIL6 inhibited N-acetyl-p-aminophenol (APAP)-induced cell death that mimicked inhibition of IL11 signaling and was unrelated to STAT3 phosphorylation. In mice, expression of HyperIL6 reduced liver damage due to IL11 dosing or APAP and promoted hepatic regeneration in a STAT3-independent manner. Following APAP, mice deleted for *Il11* were protected from liver failure and exhibited spontaneous regeneration. Despite robustly activating STAT3, HyperIL6 had no beneficial effect in *Il11* null mice. These data overturn the premise that IL6 promotes liver regeneration, show STAT3 activation to be redundant and suggest IL11 as a focus for regenerative medicine.

## Introduction

The liver has an extraordinary capacity to regenerate in response to injury. Replication of hepatocytes in midlobular zone two underlies liver regeneration (Wei et al., 2021) with a large number of cytokines and growth factors implicated as mitogens (Michalopoulos and Bhushan, 2021). Interleukin 6 (IL6) is a member of the larger IL6 family of cytokines, which include IL11 and LIF among others, and binds with high affinity to its receptor (IL6R) to signal in *cis* through a glycoprotein 130 (gp130) heterodimer via STAT3. Of all the cytokines implicated in liver regeneration, IL6 is believed to be a predominant auxiliary mitogen (Galun and Rose-John, 2013; Michalopoulos and Bhushan, 2021; Schmidt-Arras and Rose-John, 2016; Widjaja et al., 2020). This belief is anchored on a seminal study performed in mice globally deleted for *Il6*, which exhibit reduced STAT3 activity and lesser liver regeneration following injury (Cressman et al., 1996).

IL6 can bind to a soluble form of its receptor (sIL6R) to signal in *trans* to activate IL6 signaling in cells that express gp130 but low/or no IL6R (Schmidt-Arras and Rose-John, 2016). This led to the design of an artificial fusion protein of a truncated form of human IL6R linked to human IL6 (HyperIL6). HyperIL6 stimulates STAT3 signaling up to 1000-fold stronger than the respective separate molecules with a high affinity for gp130 (Fischer et al., 1997; Peters et al., 1998). The HyperIL6 superagonist can reverse fulminant liver failure due to toxin-induced liver damage (Galun et al., 2000; Hecht et al., 2001) and stimulate liver regeneration after partial hepatectomy (Peters et al., 2000). The restorative activity of HyperIL6 has also been shown for nerve (Leibinger et al., 2021) and kidney regeneration (Nechemia-Arbely et al., 2008) and in cardioprotection (Matsushita et al., 2005).

We recently found that IL11, an IL6 family protein, is hepatotoxic and prevents liver regeneration following liver injury (Dong et al., 2021; Widjaja et al., 2019, n.d.). We also showed that proteins that compete with IL11 for binding to its IL11RA/gp130 can prevent hepatotoxicity, as does genetic deletion of *Il11* (Widjaja et al., n.d.). HyperIL6 binds to gp130 in a similar way to IL11:IL11RA complexes and thus could competitively inhibit IL11 signaling. This infers that the mode of action of HyperIL6 in liver regeneration may be independent of STAT3 activity.

## Results

### STAT-independent HyperIL6 activity inhibits N-acetyl-p-aminophenol (APAP)- and IL11-induced hepatocyte cell death

To test our hypothesis, we studied APAP-induced hepatotoxicity. APAP poisoning is a common cause of liver damage, associated with impaired liver regeneration (Bernal and Wendon, 2013; Widjaja et al., n.d.). In primary human hepatocytes cultures, APAP caused approximately 40% of cell death (**Figure 1A-B; Figure 1-figure supplement 1A-B**). Inhibition of IL11 signaling using a neutralizing IL11RA antibody (X209) reduced ERK, JNK and NOX4 activity and cell death (**Figure 1A-C**). These phenotypes were mirrored by inhibition of gp130. HyperIL6 also inhibited APAP-induced cell death and this was associated with mildly increased STAT3 phosphorylation and lesser ERK, JNK and NOX4 activity (**Figure 1A-C; Figure 1-figure supplement 1A-B**).

**Figure 1.**
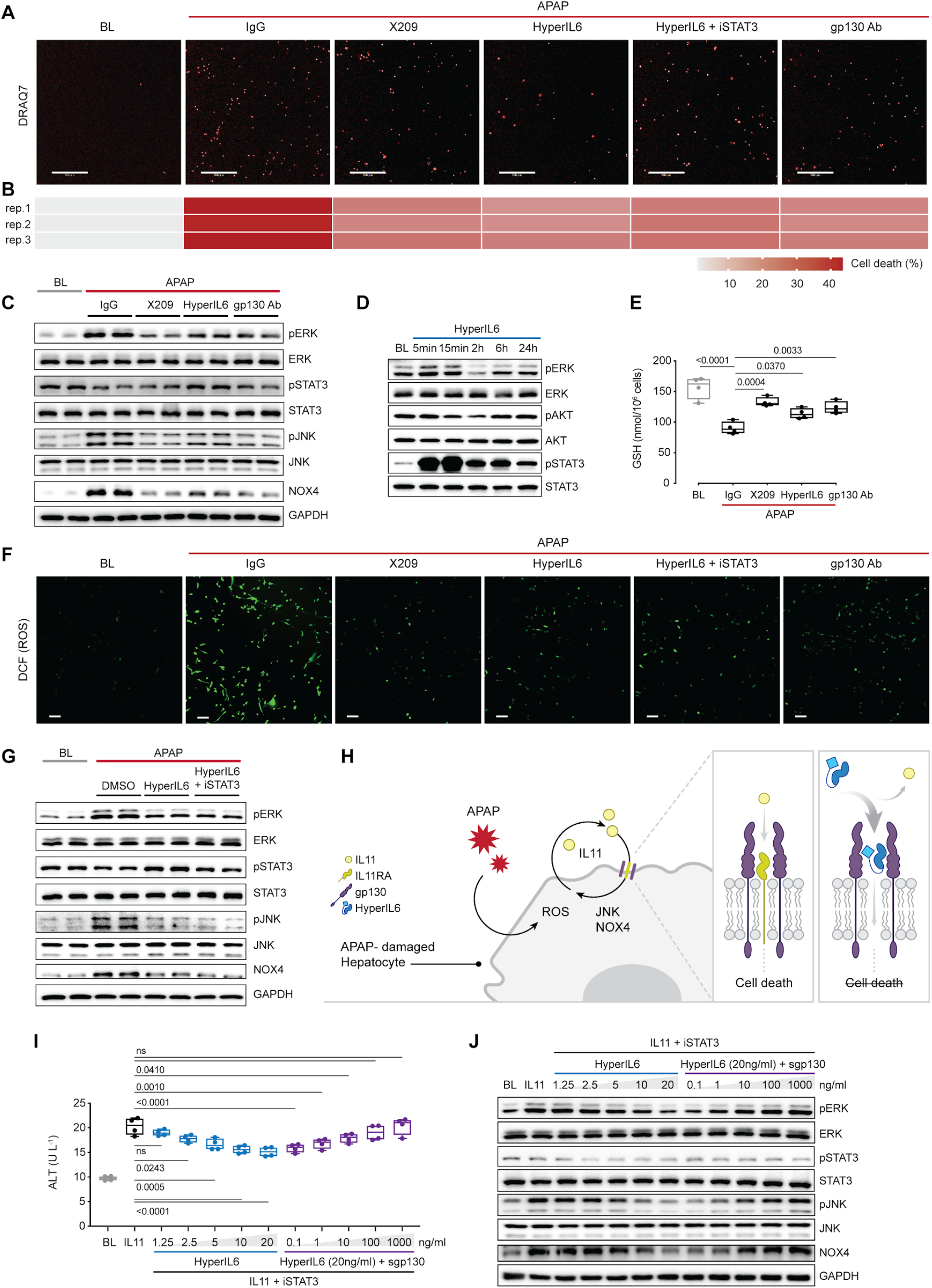
STAT-independent HyperIL6 activity inhibits APAP- or IL11-stimulated cell death through competitive binding to the shared gp130 co-receptor. (**A**) Representative fluorescent images and (**B**) quantification of DRAQ7 staining for cell death (scale bars, 200 µm) (n=3 independent experiments, 23 images per experiment) in APAP (20 mM) -treated hepatocytes in the presence of IgG (2 µg/ml), DMSO, anti-IL11RA (X209, 2 µg/ml), HyperIL6 (20 ng/ml), HyperIL6 supplemented with iSTAT3 (S3I-201, 20 µM), or anti-gp130 (2 µg/ml). (**C**) Western blots showing phospho-ERK, ERK, phospho-STAT3, STAT3, phospho-JNK, JNK, NOX4, and GAPDH levels in APAP-treated hepatocytes in the presence of IgG, X209, HyperIL6, or anti-gp130. (**D**) Western blots of phosphorylated ERK, AKT and STAT3 protein and their respective total expression in hepatocytes in response to HyperIL6 stimulation. (**E**) GSH levels (n=4) in APAP-treated hepatocytes. (**F**) Representative fluorescent images of DCFDA (2’, 7’-dichlorofluorescein diacetate) staining for ROS detection (scale bars, 100 µm) (n=4 independent experiments, 10 images per experiment) in APAP-treated hepatocytes. (**G**) Western blots showing ERK, STAT3 and JNK activation status, NOX4 protein expression in APAP-treated hepatocytes in the presence of DMSO, HyperIL6, or HyperIL6 supplemented with iSTAT3. (**H**) Proposed mechanism for competition of IL11 cis-signaling and IL6 trans-signaling by binding to gp130. (**I**) ALT secretion (n=4) and (**J**) Western blots showing ERK, STAT3, and JNK activation status, NOX4 protein expression by rhIL11 (10 ng/ml) -treated hepatocytes following a dose range stimulation of either HyperIL6 or sgp130 in the presence of iSTAT3. (**A-G, I-J**) Primary human hepatocytes; (**A-C, E-G, I-J**) 24 h stimulation. (**E, I**) Data are shown as box-and-whisker with median (middle line), 25th–75th percentiles (box) and min-max values (whiskers), one-way ANOVA with Dunnett’s correction.

HyperIL6 markedly induced STAT3 phosphorylation, had minimal effect on ERK and no effect on AKT (**Figure 1D**). Inhibition of IL11 signaling with X209 or anti-gp130 reduced APAP-induced reactive oxygen species (ROS) and maintained cellular glutathione (GSH) levels, which was also true for HyperIL6 (**Figure 1E-F; Figure 1-figure supplement 1C**). These initial studies show that HyperIL6 uniquely activates STAT3 but otherwise inhibits APAP-induced signaling and cellular phenotypes similar to neutralising IL11RA or gp130 antibodies (**Figure 1A-C, E-F; Figure 1-figure supplement 1C**).

We then examined the functional relevance of HyperIL6-induced STAT3 activation in hepatocytes exposed to APAP. S3I-201 (a STAT3 inhibitor) had no effect on the cytoprotection afforded by HyperIL6 despite inhibiting STAT3 activation. Furthermore, S3I-201 neither had any effects on HyperIL6-induced cell death, ROS induction or GSH depletion nor HyperIL6-mediated inhibition of ERK, JNK or NOX4 activity (**Figure 1A-C, E-G; Figure 1-figure supplement 1C-D**). These experiments suggest that the beneficial effects of HyperIL6 are unrelated to STAT3 activity but instead reflect a competition with IL11:IL11RA for gp130 binding, as per our hypothesis (**Figure 1H**).

We next examined whether HyperIL6 inhibited hepatotoxicity due to IL11 itself. Incubation of hepatocytes with IL11 resulted in ERK, JNK and NOX4 activation and cell death, similar to that seen with APAP (**Figure 1I-J; Figure 1-figure supplement 1E-F**). HyperIL6 dose-dependently inhibited IL11 toxicity and this was independent of STAT3 phosphorylation and could be titrated away by the addition of soluble gp130 (sgp130) (**Figure 1I-J; Figure 1-figure supplement 1E-F**). In binding assays, gp130 bound to HyperIL6 equally strongly as to an IL11:IL11RA construct (HyperIL11) (K_D_= 1 nM and 0.95 nM, respectively) whereas IL6 alone did not bind to gp130 (**Figure 1-figure supplement 2A-B**)

### Hepatocyte-specific expression of HyperIL6 prevents APAP-induced liver injury

We next studied the effects of HyperIL6 on APAP-induced liver injury (AILI) *in vivo*. Earlier studies reported the use of HyperIL6 made from human IL6 and IL6R in the mouse experiments (Galun et al., 2000; Hecht et al., 2001). This could have unappreciated off-target effects, toxicities and/or immunogenicity issues as human IL6 and IL6R have limited conservation with mouse orthologs (41% and 53.4%, respectively). Therefore, we examined the effects of recombinant mouse HyperIL6 (rm-HyperIL6) versus recombinant human HyperIL6 (rh-HyperIL6) in the mouse model of AILI (**Figure 2A**). We found that both constructs equally reduced serum ALT and AST levels and GSH depletion (**Figure 2B-D**), activated STAT3, and inhibited ERK and JNK phosphorylation (**Figure 2E**). Histology showed both constructs to reduce centrilobular necrosis, pathognomonic of APAP liver damage (**Figure 2F**).

**Figure 2.**
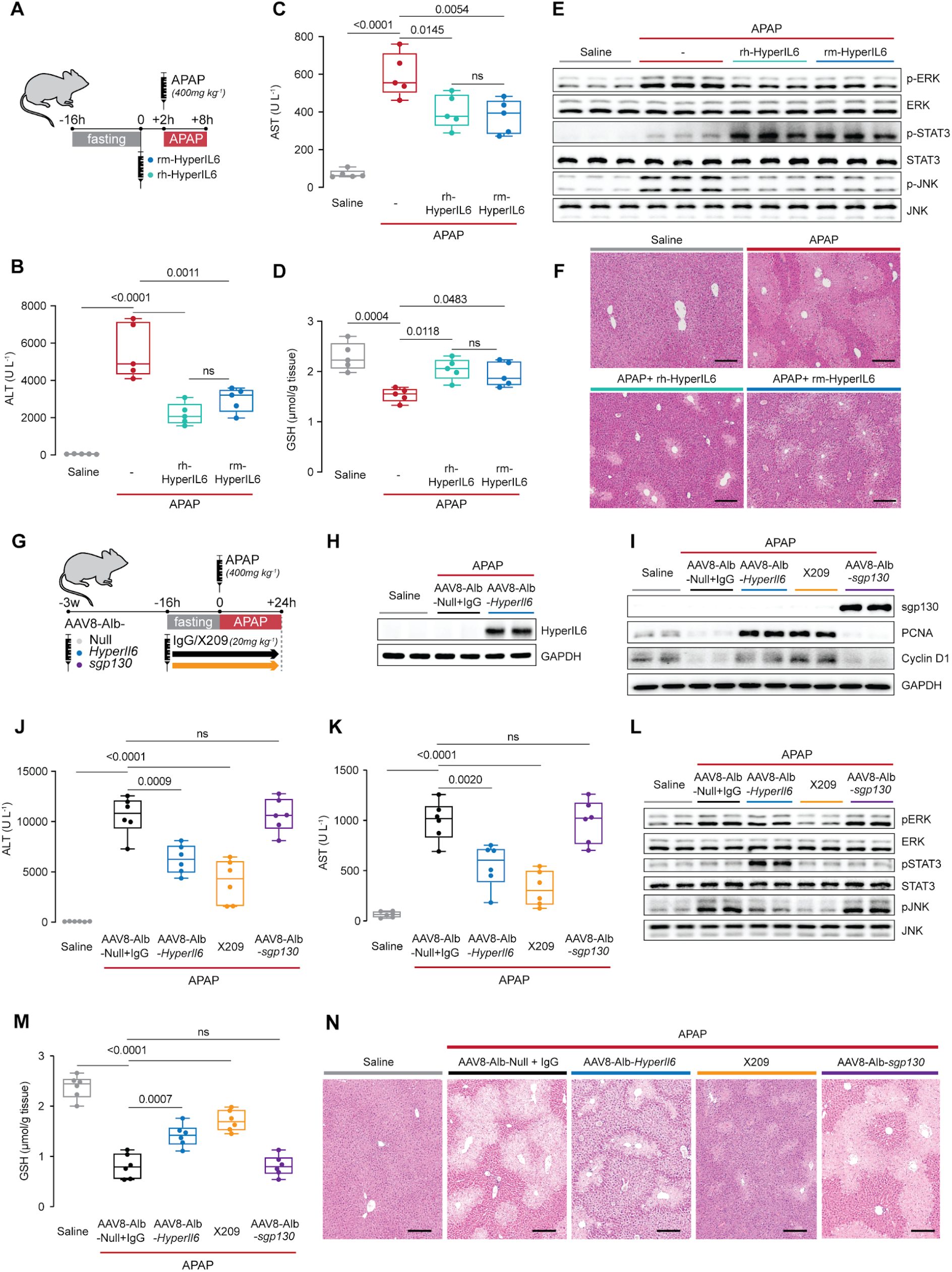
Hepatocyte-specific *HyperIl6* expression reduces APAP-induced liver injury and phenocopies the effects seen with inhibition of IL11 signaling. (**A**) Schematic of mice receiving rh-HyperIL6 or rm-HyperIL6 (500 µg/kg) administration 2 h prior to APAP injection; mice were harvested 6 h post saline or APAP injection. (**B**) Serum ALT levels, (**C**) serum AST levels, (**D**) hepatic GSH levels, (**E**) Western blot analysis of hepatic ERK, STAT3, and JNK activation, and (**F**) representative H&E-stained liver images (scale bars, 50 µm) for experiments shown in **A**. (**G**) Schematic of APAP-injected mice with hepatocyte-specific expression of *HyperIl6*/*sgp130* or IgG/X209 administration. Three weeks following AAV8-Alb-Null, AAV8-Alb-*HyperIl6*, or AAV8-Alb-*sgp130* virus injection, mice were injected with APAP (400 mg/kg); X209 or IgG (20 mg/kg) was administered at the beginning of fasting period, 16 h prior to APAP injection; control mice received saline injection; mice were harvested 24 h post saline or APAP injection. (**H**) Western blots of hepatic HyperIL6 expression and GAPDH as internal control, (**I**) Western blots showing hepatic levels of sgp130, PCNA, Cyclin D1, and GAPDH as internal control, (**J**) serum ALT levels, (**K**) serum AST levels, (**L**) Western blots showing hepatic levels of phospho-ERK, ERK, phospho-STAT3, STAT3, phospho-JNK, and JNK, (**M**) hepatic GSH levels, and (**N**) representative H&E-stained liver images (scale bars, 50 µm) for experiments shown in **G**. (**B-D**) N=5 mice/group; (**J-K, M**) n=6 mice/group. (**B-D, J-K, M**) Data are shown as box-and-whisker with median (middle line), 25th–75th percentiles (box) and min-max values (whiskers), one-way ANOVA with Tukey’s correction.

We therefore used species-matched mouse HyperIL6 for hepatocyte-specific *HyperIl6* expression studies. Mice were injected with adeno-associated virus serotype 8 (AAV8) encoding either albumin promoter-driven mouse *HyperIl6* (AAV8-Alb-*HyperIl6*) or one of two controls: AAV8-Alb-*sgp130* or AAV8-Alb-Null. AAV8-Alb-*sgp130*, which encodes mouse sgp130, provides a second viral control group while probing for effects of endogenous IL6 *trans*-signaling. We compared data from the AAV8-treated mice with a group where we inhibited IL11 signaling by X209 (**Figure 2G**).

The day after APAP, mice over-expressing HyperIL6 (**Figure 2H**) had lower ALT/AST levels as compared to AAV8-Alb-Null group (**Figure 2J-K**). Mice receiving X209 or HyperIL6 were protected from liver damage as compared to the AAV8-Alb-Null+IgG control group (**Figure 2J-K**). AAV8-Alb-*sgp130* induced high sgp130 expression but had no effect on APAP-induced liver injury, ruling out an endogenous protective *trans* activity for IL6 complexes (**Figure 2I-K**).

Liver regeneration is associated with a signature of increased PCNA and Cyclin D1 expression (Sekiya and Suzuki, 2011), which was apparent 24 h post APAP in both HyperIL6 expressing or X209-treated mice but not in AAV8-Alb-Null+IgG or in sgp130 groups (**Figure 2I**). HyperIL6 or X209 partially restored liver GSH levels and inhibited ERK and JNK activation whereas STAT3 was uniquely activated in HyperIL6-expressing mice (**Figure 2L-M**). Histology showed typical centrilobular necrosis in APAP-treated AAV8-Alb-Null or sgp130 mice, which was lesser in mice expressing HyperIL6 or following X209 administration (**Figure 2N**).

Taken together, these data show that both human and mouse HyperIL6 are protective against APAP liver damage in mice and that hepatocyte-specific HyperIL6 expression is protective and promotes liver regeneration. However, the beneficial effects of HyperIL6 appear unrelated to STAT3 activation and instead mirror precisely the protection seen with inhibition of IL11-driven ERK/JNK/NOX4 activation.

### The protective effects of HyperIL6 in the liver are STAT3-independent

To formally exclude a protective role for STAT3 signaling in the liver, we studied the effects of HyperIL6 in the context of S3I-201 (10 mg/kg) administration prior to AILI (**Figure 3A**). Following APAP, mice with hepatocyte-specific *HyperIl6* expression had reduced serum ALT/AST levels, restored hepatic GSH levels, lesser ERK and JNK activity along with diminished centrilobular necrosis (**Figure 3B-F**). We observed elevated STAT3 phosphorylation in APAP-treated (6 h) control mice that was further increased in AAV8-Alb-*HyperIl6* mice but absent in mice receiving S3I-201 (**Figure 3E**). The beneficial effects of HyperIL6 on the liver were unaffected by STAT3 inhibition.

**Figure 3.**
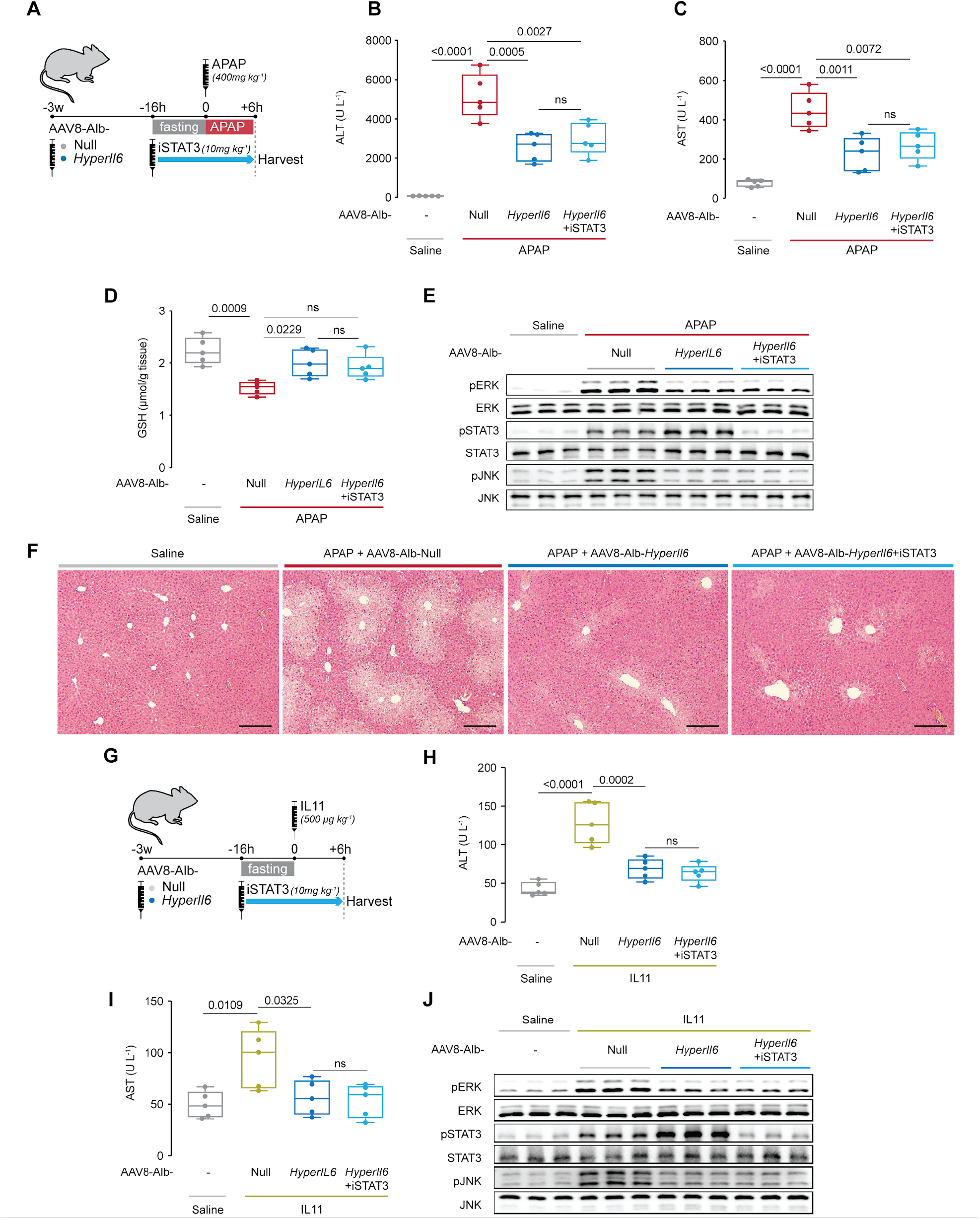
Hepatocyte-specific *HyperIl6* expression reduces APAP- or IL11-induced liver injury independent of STAT3 activation. (**A**) Schematic of APAP-injected mice with hepatocyte-specific expression of *HyperIl6* ± iSTAT3 administration. Three weeks following AAV8-Alb-Null or AAV8-Alb-*HyperIl6* virus injection, mice were injected with APAP (400 mg/kg); iSTAT3 (S3I-201, 10 mg/kg) was administered at the beginning of fasting period, 16 h prior to APAP injection; control mice received saline injection; mice were harvested 6 h post saline or APAP injection. (**B**) Serum ALT levels, (**C**) serum AST levels, (**D**) hepatic GSH levels, (**E**) Western blots showing hepatic phospho-ERK, ERK, phospho-STAT3, STAT3, phospho-JNK, and JNK, and (**F**) representative H&E-stained liver images (scale bars, 50 µm) for experiments shown in **A**. (**G**) Schematic of rmIL11-injected mice with hepatocyte-specific expression of *HyperIl6* ± iSTAT3 administration. Mice were injected with rmIL11 (500 µg/kg), three weeks following AAV8-Alb-Null or AAV8-Alb-*HyperIl6* virus injection; iSTAT3 (10 mg/kg) was administered at the beginning of fasting period, 16 h prior to rmIL11 injection; control mice received saline injection; mice were harvested 6 h post saline or IL11 injection. (**H**) Serum ALT levels, (**I**) serum AST levels, and (**J**) Western blots showing hepatic ERK, STAT3 and JNK activation status for experiments shown in **G**. (**B-D, H-I**) N=5 mice/group; data are shown as box-and-whisker with median (middle line), 25th–75th percentiles (box) and min-max values (whiskers), one-way ANOVA with Tukey’s correction.

Our hypothesis (**Figure 1H**) and data (**Figures 1 and 2**) suggest that the beneficial effects of HyperIL6 could be related to its competitive inhibition of IL11. To test this directly, we injected recombinant mouse IL11 (rmIL11) to mice with *HyperIl6* expression with or without administration of S3I-201 (**Figure 3G**). Injection of rmIL11 to mice resulted in elevated ALT/AST levels and activation of ERK and JNK (**Figure 3H-J**). Mice expressing HyperIL6 exhibited STAT3 phosphorylation, had lower IL11-induced ALT/AST levels and lesser activation of ERK and JNK. Notably, administration of S3I-201 to mice expressing HyperIL6 reduced phospho-STAT3 levels to baseline but did not diminish its beneficial effects at any level of assessment. (**Figure 3H-J**).

### HyperIL6 has no effect on APAP-induced liver injury in mice deleted for *Il11*

If the protective effects of HyperIL6 are due to its inhibition of IL11 signaling then HyperIL6 should be ineffective in APAP injury in the absence of IL11. Thus we studied the impact of HyperIL6 on AILI in mice globally deleted for *Il11* (*Il11*^*-/-*^) (**Figure 4A**).

**Figure 4.**
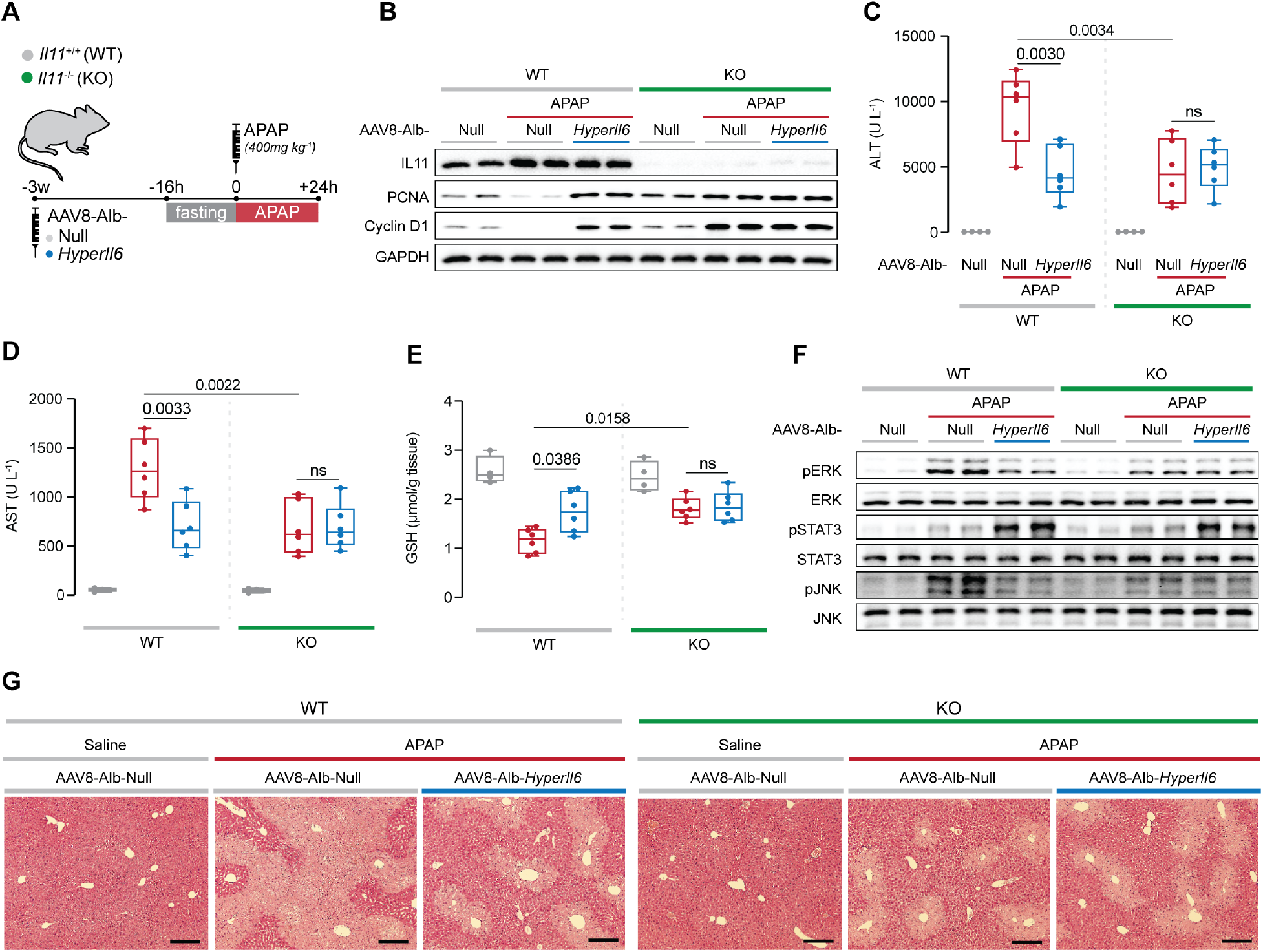
Hepatocyte-specific *HyperIl6* expression has no benefit in *Il11*^*-/-*^ mice. (**A**) Schematic of APAP injury in *Il11*^*-/-*^ and *Il11*^*+/+*^ mice (control) with hepatocyte-specific expression of *HyperIl6*. Three weeks following AAV8-Alb-Null or AAV8-Alb-*HyperIl6* virus injection, overnight-fasted *Il11*^*+/+*^ and *Il11*^*-/-*^ mice were injected with saline or APAP (400 mg/kg); mice were harvested 24 h post saline or APAP injection. (**B**) Serum ALT levels. (**C**) Serum AST levels. (**D**) Hepatic GSH levels. (**E**) Western blots showing hepatic phospho-ERK, ERK, phospho-STAT3, STAT3, phospho-JNK, and JNK. (**F**) Representative H&E-stained liver images (scale bars, 50 µm). (**C-E**) Saline (n=4 mice/group), APAP (n=6 mice/group); data are shown as box-and-whisker with median (middle line), 25th–75th percentiles (box) and min-max values (whiskers), 2-way ANOVA with Sidak’s correction.

APAP dosing resulted in increased IL11 expression in the injured livers of wildtype (WT) mice that was absent in *Il11*^*-/-*^ mice (**Figure 4B**). As compared to WT controls, expression of HyperIL6 in WT mice was associated with lesser liver damage and a molecular signature of regeneration following APAP administration (**Figure 4B-E, G**). Following APAP and as compared to WT mice, *Il11*^*-/-*^ mice had reduced ALT, AST, and centrilobular necrosis, higher GSH levels along with increased PCNA and Cyclin D1 expression but these phenotypes were unaffected by HyperIL6 expression (**Figure 4B-E, G**). Thus, expression of HyperIL6 in *Il11*^*-/-*^ mice had no additive protective effect on liver damage or on liver regeneration.

At the signaling level, APAP-induced ERK and JNK activation were reduced in both HyperIL6-expressing WT mice and in *Il11*^*-/-*^ mice, which correlated with beneficial effects (**Figure 4B-G**). HyperIL6 expression robustly increased STAT3 phosphorylation in both WT and *Il11*^*-/-*^ mice but this activity was unrelated to liver protection or regeneration (**Figure 4B-G**).

## Conclusion

For two decades IL6 signaling, and in particular HyperIL6 activation of STAT, has been thought to promote regeneration. While some early reports questioned this assertion (Sakamoto et al., 1999), it is now generally accepted (Schmidt-Arras and Rose-John, 2016). Here we show that HyperIL6-mediated inhibition of IL11 signaling in injured hepatocytes, latent until now, underlies the hepatoprotective effects of HyperIL6. In keeping with this the protective effects of HyperIL6 are STAT3-independent. This challenges the interpretation of a body of literature and suggests IL11 instead of IL6 as a focus for regenerative studies of the liver (Nechemia-Arbely et al., 2011) and perhaps nerves (Leibinger et al., 2021) and kidney (Nechemia-Arbely et al., 2008).

## Materials and methods

### Ethics statements

All experimental protocols involving human subjects (commercial primary human cell lines) have been performed in accordance with the ICH Guidelines for Good Clinical Practice. As written in the datasheets, ethical approval of primary human hepatocytes has been obtained by ScienCell.

Animal studies were carried out in compliance with the recommendations in the Guidelines on the Care and Use of Animals for Scientific Purposes of the National Advisory Committee for Laboratory Animal Research (NACLAR). All experimental procedures were approved (SHS/2014/0925 and SHS/2019/1482) and conducted in accordance with the SingHealth Institutional Animal Care and Use Committee.

### Antibodies

phospho-AKT (4060, CST), AKT (4691, CST), Cyclin D1 (55506, CST), phospho-ERK1/2 (4370, CST), ERK1/2 (4695, CST), GAPDH (2118, CST), gp130 (extracellular, PA5-77476, Thermo Fisher), gp130 (neutralizing, MAB628, R&D systems), IgG (Aldevron), IL6 (AF506, R&D systems), p-JNK (4668, CST), JNK (9252, CST), IL11 (X203, Aldevron), IL11RA (neutralizing, X209, Aldevron), NOX4 (MA5-32090, Invitrogen), PCNA (13110, CST), phospho-STAT3 (4113, CST), STAT3 (4904, CST), anti-mouse HRP (7076, CST), anti-rabbit HRP (7074, CST), anti-rat HRP (ab97057, Abcam).

### Recombinant proteins

Commercial recombinant proteins: rhIL11 (UniProtKB: P20809, Genscript), rmIL11 (UniProtKB: P47873, Genscript), rh-HyperIL6 (IL6R:IL6 fusion protein, 8954-SR, R&D systems), rm-HyperIL6 (IL6R:IL6 fusion protein, 9038-SR, R&D systems), human soluble gp130 Fc (671-GP-100, R&D systems).

### Chemical

APAP (A3035, Sigma), iSTAT3 (S3I-201, SML0330, Sigma).

### AAV8 vectors

All AAV8 vectors used in this study were synthesized by Vector Biolabs. AAV8 vector carrying mouse *HyperIl6* cDNA driven by Albumin (Alb) promoter is referred to as *AAV8-Alb-HyperIl6*, which was constructed using the cDNA sequences of mouse IL-6/IL-6R alpha fusion protein (9038-SR, R&D systems). AAV8-Null vector was used as vector control.

### Cell culture

Primary human hepatocytes (5200, ScienCell) were maintained in hepatocyte medium (5201, ScienCell) supplemented with 2% fetal bovine serum, 1% Penicillin-streptomycin at 37°C and 5% CO_2_. Hepatocytes were serum-starved overnight unless otherwise specified in the methods prior to 24 h stimulation with different doses of various recombinant proteins as outlined in the main text and/or figure legends. All experiments were carried out at low cell passage (<P3).

### Operetta high throughput phenotyping assay

Primary human hepatocytes were seeded in 96-well black CellCarrier plates (PerkinElmer) at a density of 5×10^3^ cells per well. Following stimulations, cells were incubated 1 h with 1µg/ml Hoechst 33342 (62249, Thermo Fisher Scientific) and DRAQ7 (D15106, Thermo Fisher Scientific) in serum-free basal medium. Each condition was imaged from triplicated wells and a minimum of 23 fields/well using Operetta high-content imaging system 1483 (PerkinElmer). Live and dead cells were quantified using Harmony v3.5.2 (PerkinElmer).

### Reactive Oxygen Species (ROS) Detection

Primary human hepatocytes were seeded on 8-well chamber slides (1.5×10^4^ cells/well). For this experiment, cells were not serum-starved prior to treatment. 24 h following stimulation, cells were washed, incubated with 25 µM of DCFDA solution (ab113851, abcam) for 45 min at 37°C in the dark, and rinsed with the dilution buffer according to the manufacturer’s protocol. Live cells with positive DCF staining were imaged with a filter set appropriate for fluorescein (FITC) using a fluorescence microscope (Leica).

### Animal models

Animal procedures were approved and conducted in accordance with the SingHealth Institutional Animal Care and Use Committee (IACUC). All mice were housed in temperatures of 21-24°C with 40-70% humidity on a 12 h light/12 h dark cycle and provided food and water ad libitum, except in the fasting period, during which only water was provided ad libitum.

### Mouse models of Acetaminophen (APAP)

Prior to APAP, 9-12 weeks old male mice were fasted overnight. Mice were given APAP (400 mg/kg) by intraperitoneal (IP) administration and euthanized 6 h or 24 h post APAP, as outlined in the main text or figure legends.

### In vivo administration of rh-HyperIL6, rm-HyperIL6, or rmIL11

rh-HyperIL6, rm-HyperIL6, or rmIL11 were administered *via* IP injection at a concentration of 500 µg/kg.

### In vivo expression of HyperIL6 or sgp130

6-8-week old male C57BL/6NTac mice (InVivos, Singapore) were injected with 4×10^11^ gc AAV8-Alb-*HyperIl6* or AAV8-Alb-*sgp130* virus to induce hepatocyte specific expression of HyperIL6 or sgp130; control mice were injected with 4×10^11^ gc AAV8-Alb-Null virus. 3 weeks following virus administration, mice were given IP administration of APAP and euthanized at the time point outlined in the main text or figure legends.

### In vivo administration of anti-IL11RA (X209) or iSTAT3 (S3I-201)

C57BL/6NTac male mice were IP administered anti-IL11RA (X209, 20 mg/kg), IgG isotype control (11E10, 20 mg/kg), or iSTAT3 (S3I-201, 10 mg/kg) at the beginning of fasting period.

### Il11^−/−^mice

Mice lacking functional alleles for Il11 (*Il11*^−*/*−^), in which Crispr/Cas9 technique was used to knock out the *Il11* gene (ENSMUST00000094892.11) were generated and validated previously. 6-8-week old male *Il11*^−*/*−^ mice and their WT littermates (*Il11*^*+/+*^) were injected with 4×10^11^ gc AAV8-Alb-*HyperIl6* virus to induce hepatocyte specific expression of HyperIL6; control mice were injected with 4×10^11^ gc AAV8-Alb-Null virus. 3 weeks following virus administration, mice were given IP administration of APAP and euthanized 24 h post APAP.

### Colorimetric assays

The levels of alanine transaminase (ALT) or aspartate aminotransferase (AST) in mouse serum and hepatocyte supernatant were measured using ALT Activity (ab105134, Abcam) or AST (ab105135, Abcam) Assay Kits. Liver glutathione sulfhydryl (GSH) measurements were performed using Glutathione Colorimetric Detection Kit (EIAGSHC, Thermo Fisher). All colorimetric assays were performed according to the manufacturer’s protocol.

### Immunoblotting

Western blots were carried out from hepatocyte and liver tissue lysates. Hepatocytes and tissues were homogenized in radioimmunoprecipitation assay (RIPA) buffer containing protease and phosphatase inhibitors (Thermo Fisher), followed by centrifugation to clear the lysate. Protein concentrations were determined by Bradford assay (Bio-Rad). Equal amounts of protein lysates were separated by SDS-PAGE, transferred to PVDF membrane, and subjected to immunoblot analysis for the indicated primary antibodies. Proteins were visualized using the ECL detection system (Pierce) with the appropriate secondary antibodies.

### Surface plasmon resonance (SPR)

SPR measurements were performed on a BIAcore T200 (GE Healthcare) at 25°C. Buffers were degassed and filter-sterilized through 0.2 μm filters prior to use. Gp130 was immobilized onto a carboxymethylated dextran (CM5) sensor chip using standard amine coupling chemistry. For kinetic analysis, a concentration series (0.39 nM to 120 nM) of IL6, HyperIL11 or HyperIL6 was injected over the gp130 and reference surfaces at a flow rate of 30 μl/min. All the analytes were dissolved in HBS-EP+ (BR100669, GE Healthcare) containing 1 mg/ml BSA. The association and dissociation were measured for 210s and 300s respectively. After each analyte injection, the surface was regenerated by two times injection of Glycine-HCl (10mM, pH1.5), followed by a 5 min stabilisation period. All sensorgrams were aligned and double-referenced. Affinity and kinetic constants were determined by fitting the corrected sensorgrams with the 1:1 Langmuir model using BIAevaluation v3.0 software (GE Healthcare). The equilibrium binding constant *K*_*D*_ was determined by the ratio of the binding rate constants *k*_*d*_*/k*_*a*_.

### Hematoxylin & Eosin (H&E) staining

Livers were fixed for 48 h at room temperature (RT) in 10% neutral-buffered formalin (NBF), dehydrated, embedded in paraffin blocks and sectioned at 7 μm. Sections were stained with H&E according to standard protocol and examined by light microscopy.

### Statistical analysis

Statistical analyses were performed using GraphPad Prism software (version 6.07). For comparisons between more than two conditions, one-way ANOVA with Dunnett’s correction (when several conditions were compared to one condition) or Tukey’s correction (when several conditions were compared to each other) were used. Comparison analysis for several conditions from two different groups were performed by 2-way ANOVA and corrected with Sidak’s multiple comparisons when the means were compared to each other. The criterion for statistical significance was P < 0.05.

## Acknowledgements

This research was supported by the National Medical Research Council (NMRC), Singapore STaR awards (NMRC/STaR/0029/2017), NMRC Centre Grant to the NHCS, MOH-CIRG18nov-0002, MRC-LMS (UK), Goh Foundation, Tanoto Foundation to S.A.C. A.A.W. is supported by NMRC/OFYIRG/0053/2017. The authors would like to acknowledge the technical support of B.L.George and J.Tan.

## Supplementary figures

**Figure 1-figure supplement 1.**
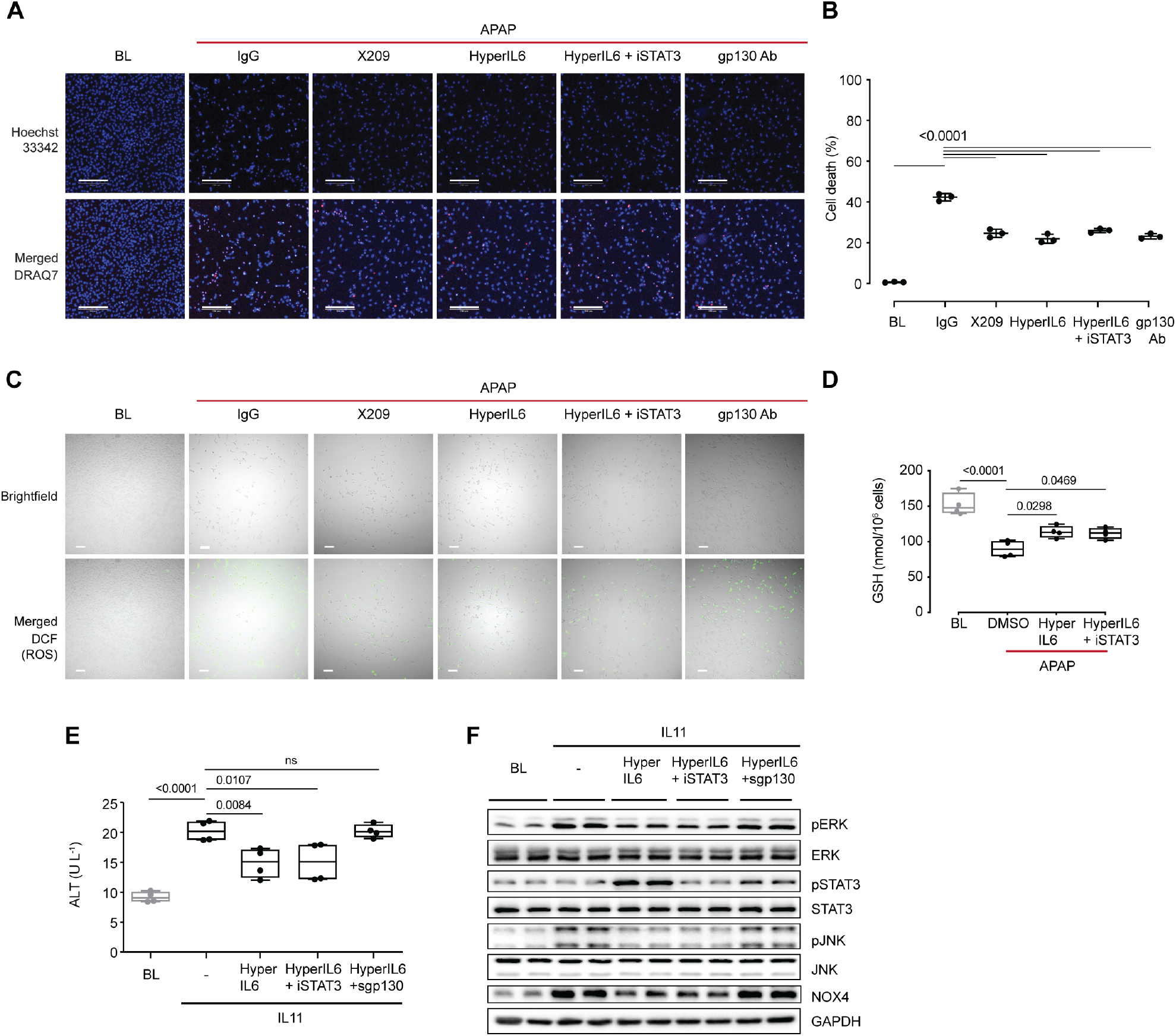
STAT-independent HyperIL6 activity inhibits IL11-stimulated hepatocyte cell death. (**A**) Representative Hoechst 33342 and merged fluorescence images (scale bars, 200 µm) and (**B**) quantification of cell death for DRAQ7 staining experiment (n=3 independent experiments, 23 images per experiment) shown in **Figure 1A-B**. (**C**) Representative brightfield images and merged fluorescence-brightfield images (scale bars, 100 µm) for DCFDA staining experiment (n=4 independent experiments, 10 images per experiment) shown in **Figure 1F**. (**D**) GSH levels (n=4) in APAP-treated hepatocytes. (**E**) ALT secretion (n=4) and (**F**) Western blots showing ERK, STAT3, and JNK activation status, NOX4 protein expression in rhIL11 (10 ng/ml) -treated hepatocytes in the presence of HyperIL6 (20 ng/ml), HyperIL6 supplemented with iSTAT3 (S3I-201, 20 µM), or sgp130 (1 µg/ml). (**B, D, E**) Data are shown as box-and-whisker with median (middle line), 25th–75th percentiles (box) and min-max values (whiskers), one-way ANOVA with Dunnett’s correction.

**Figure 1-figure supplement 2.**
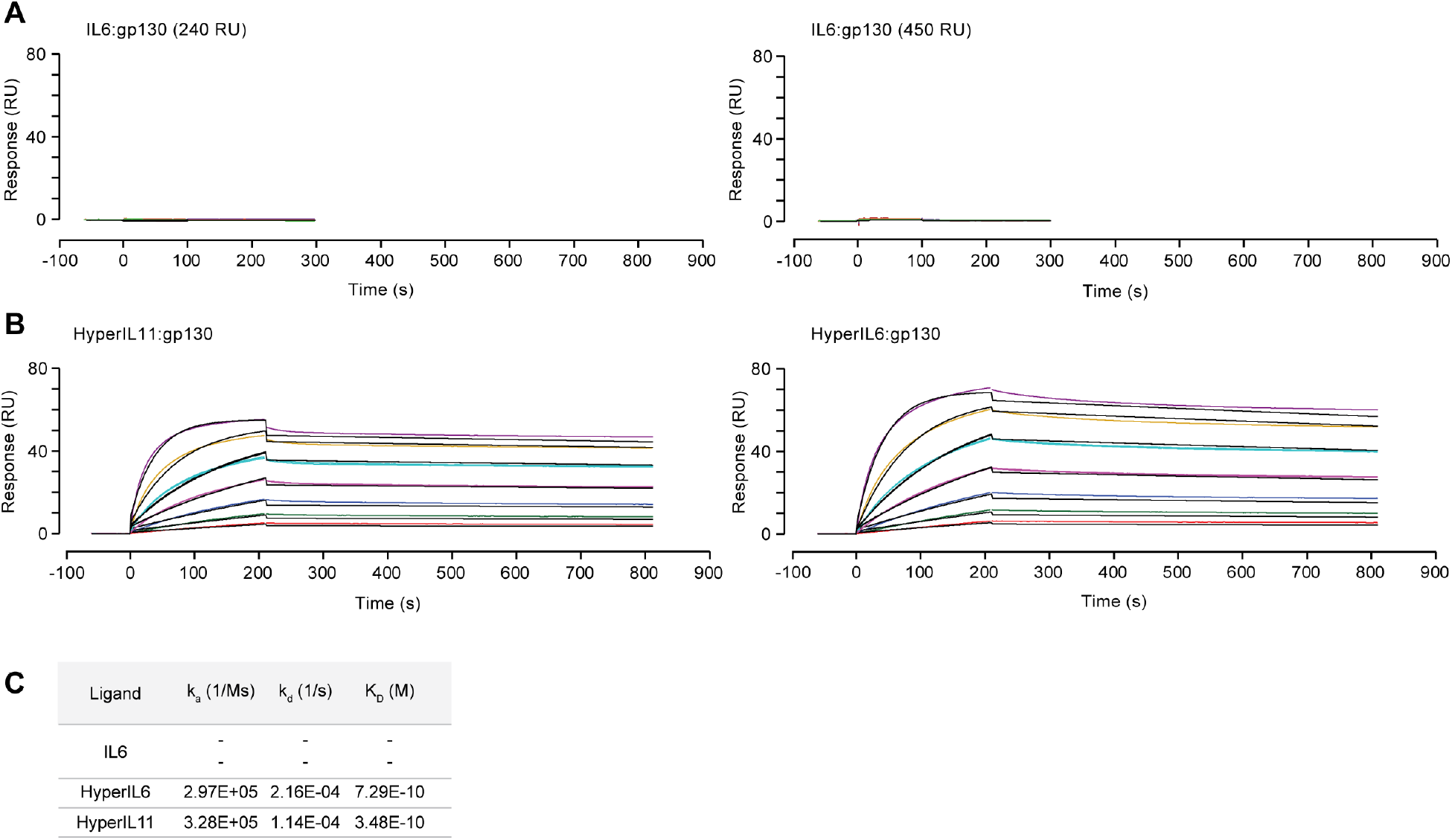
Surface plasmon resonance analysis of IL6, HyperIL11, or HyperIL6 binding to gp130. Sensorgrams showing binding of (**A**) IL6, (**B**) HyperIL11 (left) and HyperIL6 (right) to immobilized gp130. The colored lines represent the experimental data; the black lines represent a theoretically fitted curve (1:1 Langmuir). (C) Binding affinity, association and dissociation constants for experiments shown in **A and B**.

